# Differential control of corticotroph Ca^2+^ signalling by corticotrophin-releasing hormone and arginine vasopressin

**DOI:** 10.64898/2026.05.07.723482

**Authors:** Sacha M James, Isabella Marinelli, Thomas Pons, Nicola Romanò, Joël Tabak, Pauline Campos, Jamie J Walker

**Affiliations:** Department of Mathematics and Statistics, Faculty of Environment, Science and Economy, University of Exeter, Exeter, United Kingdom; College of Medicine and Health, University of Birmingham, Birmingham, United Kingdom; Department of Natural Sciences, Faculty of Environment, Science and Economy, University of Exeter, Exeter, United Kingdom; Institute for Neuroscience and Cardiovascular Research, Edinburgh Medical School, University of Edinburgh, United Kingdom; ZJU-UoE Institute, Zhejiang University, School of Medicine, Haining, China; Department of Clinical and Biomedical Science, Faculty of Health and Life Sciences, University of Exeter, Exeter, United Kingdom

**Keywords:** HPA axis, corticotroph, stress, CRH, AVP

## Abstract

Corticotroph cells convert hypothalamic inputs into adrenocorticotrophic hormone secretion via intracellular calcium (Ca^2+^) signalling, but how they integrate corticotrophin-releasing hormone (CRH) and arginine vasopressin (AVP) across physiological concentration ranges remains unclear. Here, we quantified intracellular Ca^2+^ responses in isolated rat corticotrophs to CRH and AVP, applied alone and in combination, to characterise response magnitude, temporal dynamics, and cell recruitment. Both secretagogues increased Ca^2+^ signalling in a concentration-dependent manner, but with distinct effects: AVP generally evoked larger responses, faster response onset, and greater cell recruitment than CRH when applied alone. Under co-stimulation, increasing CRH concentration increased the proportion of cells classified as synergistic without altering positive synergy values, suggesting CRH-dependent control of interaction likelihood rather than interaction strength. Marked cell-to-cell heterogeneity was observed across all conditions, consistent with corticotroph subpopulations differing in activation thresholds. Together, these findings show that AVP drives broad corticotroph recruitment, whereas CRH modulates how corticotrophs integrate convergent inputs, defining complementary roles in shaping pituitary output.

## Introduction

The hypothalamic-pituitary-adrenal (HPA) axis is the primary neuroendocrine system responsible for coordinating the physiological response to stress by regulating glucocorticoid hormone release from the adrenal glands. Glucocorticoids play a crucial role in regulating numerous physiological processes, including energy metabolism, immune function, and circadian rhythmicity [1, 2]. Glucocorticoid production is controlled by adrenocorticotrophic hormone (ACTH), which is secreted by corticotroph cells in the anterior pituitary in response to upstream stimulation from the hypothalamic secretagogues corticotrophin-releasing hormone (CRH) and arginine vasopressin (AVP). These neuropeptides are co-expressed by parvocellular neurons of the hypothalamic paraventricular nucleus (PVN) and are released from nerve terminals in the median eminence into the hypophyseal portal vasculature, through which they are transported to the anterior pituitary [3]. In the anterior pituitary, CRH and AVP bind to their respective receptors on corticotroph cells, increasing electrical activity and intracellular calcium (Ca^2+^) signalling, which culminates in the exocytosis of ACTH-containing vesicles.

Despite their shared role in stimulating corticotroph activity, CRH and AVP engage distinct intracellular signalling pathways within corticotroph cells. CRH binds to CRHR1 receptors to activate the cAMP/PKA pathway and drive Ca^2+^ influx from the extracellular environment via L-type voltage-gated calcium channels [4, 5, 6]. Activation of this pathway is characterised electrically by cell depolarisation, which results in increased spiking frequency and a transition from spiking to bursting behaviour [7]. In contrast, AVP binds to V1b receptors, stimulating the phospholipase C/PKC pathway, which increases intracellular Ca^2+^ through mobilisation of Ca^2+^ stored in the endoplasmic reticulum (ER) [8]. Electrophysiologically, AVP evokes an early transient hyperpolarisation associated with ER Ca^2+^ release, followed by a later increase in spiking activity [7, 9].

CRH and AVP play differential roles in regulating HPA axis dynamics. Under basal conditions, where HPA activity is characterised by ultradian rhythms [10, 11], pharmacological blockade of V1b receptors does not alter the ultradian pattern of corticosterone secretion [12], suggesting that basal rhythmicity is not strongly dependent on AVP signalling and may rely more heavily on CRH-dependent regulation. In response to acute stress, both CRH and AVP gene expression in the PVN are upregulated [13, 14, 15], accompanied by increased secretion of both secretagogues. During chronic or repeated stress, however, the expression and regulation of CRH and AVP become more complex. Within the first 8 days of chronic stress exposure, CRH expression is upregulated [16, 17, 18]; over longer periods of chronic stress, CRH expression may remain unchanged compared with pre-stress levels or decline [19, 15]. By contrast, AVP expression is upregulated in response to chronic stress over both shorter [20] and longer [15] exposure periods. Thus, changes in the relative levels of CRH and AVP may provide a mechanism by which hypothalamic PVN neurons encode stress-related information and tune downstream corticotroph signalling.

Given the importance of CRH–AVP balance in regulating HPA axis activity, we sought to determine how these secretagogues interact at the level of corticotroph Ca^2+^ signalling. In a previous study, we characterised heterogeneous intracellular Ca^2+^ responses of corticotroph cells to fixed physiological concentrations of CRH and AVP [21]. Building on that work, the aim of this study was to investigate how shifting the relative balance of CRH and AVP concentrations regulates corticotroph Ca^2+^ response dynamics. To address this, we performed real-time Ca^2+^ imaging in primary rat corticotroph cells and characterised their responses to a range of CRH and AVP concentrations, applied individually and in combination. Specifically, we examined how these secretagogues affected Ca^2+^ response magnitude and delay, cell recruitment, and temporal Ca^2+^ response dynamics, and quantified how co-stimulation with CRH and AVP modulated synergistic and anti-synergistic Ca^2+^ responses.

## Methods

### Animals

Male Sprague Dawley rats (Envigo RMS (UK) Ltd.) aged 8–16 weeks were used for all experiments. Animals were housed under a 12:12 h light/dark cycle with food and water provided ad libitum. All procedures complied with the UK Animal (Scientific Procedures) Act 1986 and were approved by the University of Exeter Animal Welfare and Ethical Review Body.

### Primary pituitary cell culture

Tissue collection was performed between 09:00–11:00 h, with care taken to minimise environmental stress prior to anaesthesia. Rats were anaesthetised with isoflurane and decapitated, and the pituitary gland was rapidly excised into DMEM ([+] 4.5 g/l glucose, [+] L-glutamine, [+] 25 mM HEPES, [-] pyruvate). The anterior lobe was isolated from the intermediate and posterior zones, avoiding contamination with POMC-expressing melanotrophs [22], and roughly chopped. Tissue was transferred to 2.5 ml chilled digestion medium (DMEM, trypsin (207 TAME Units/ml), DNase I (36 Kunitz Units/ml)) and incubated for 30 min at 37 °C with frequent trituration. Digestion was terminated by addition of 5 ml blocking solution (DMEM, soybean trypsin inhibitor (0.25 mg/ml), DNase I (36 Kunitz Units/ml), aprotinin (100 Kallikrein Units/ml)). The cell suspension was filtered (70 µm), centrifuged (10 min,100*×* g), resuspended in RBC lysis buffer (37 °C; Invitrogen, 00-4333-57), and centrifuged again (10 min, 100*×* g). Cells were then resuspended in growth medium containing antibiotics and antimycotics (DMEM, bovine serum albumin (BSA) solution (30% in DPBS; 1% v/v), insulin–transferrin–selenium (1% v/v), fibronectin (from bovine plasma; 0.42% v/v), antibiotic–antimycotic solution (1% v/v)). Cells (25 µl) were plated onto coverslips and incubated for 30 min at 37 °C to allow attachment, then topped up to 1 ml of this medium and cultured overnight at 37 °C.

### Lentiviral transduction

Twenty-four hours after plating, cells were transduced with a lentiviral construct encoding mCherry under the control of a minimal POMC promoter (VectorBuilder, 2.17 *×*10^9^ TU/ml) [23]. Growth medium containing antibiotics and antimycotics was replaced with antibiotic-free growth medium (DMEM, BSA solution (30% in DPBS; 1% v/v), insulin–transferrin–selenium (1% v/v), fibronectin (from bovine plasma; 0.42% v/v)). Lentivirus (1 µl; 300,000 TU) was added to each well, and cells were incubated at 37 °C for 6 h before virus removal. Cells were then incubated for a further 36 h at 37 °C in fresh antibiotic-free growth medium before fixation for immunostaining or calcium imaging.

### Immunostaining

Cells were fixed in 4 % formaldehyde (paraformaldehyde in phosphate-buffered saline (PBS)) for 1 h and washed three times with PBS. Cells were permeabilised in PBS containing 0.3 % Triton X-100 for 10 min, then blocked in PBS containing 0.05 % Triton X-100 and 0.3 % BSA for 60 min.

Cells were incubated with primary antibody solution (blocking buffer containing polyclonal rabbit anti-ACTH antisera (Invitrogen, AB_2806982; 1:1000) and 2 % normal goat serum) for 90 min with gentle agitation, then washed three times with PBS. Cells were then incubated with secondary antibody solution (blocking buffer containing goat anti-rabbit Alexa Fluor 488 (Invitrogen, AB_143165; 1:500)) for 60 min with gentle agitation, followed by three PBS washes. Coverslips were mounted using DAPI Fluoromount-G (SouthernBiotech, 0100-20) and imaged by fluorescence microscopy.

To quantify viral transduction efficiency and specificity, images were acquired in FITC and TRITC channels corresponding to ACTH immunostaining and mCherry fluorescence, respectively. Negative controls included (i) cells processed without virus and (ii) cells processed without primary antibody. Thresholds were defined from control images as the mean signal within manually defined ROIs plus two standard deviations. ACTH-positive ROIs were identified by thresholding, and mean mCherry intensity was measured within these regions. Cells were classified as co-localised if both ACTH and mCherry signals exceeded their respective thresholds. Efficiency was calculated as the proportion of ACTH-positive cells that were mCherry-positive across 12 coverslips from three independent preparations. Specificity was assessed by identifying mCherry-positive ROIs and measuring ACTH signal within these regions, and defined as the proportion of mCherry-positive cells that were also ACTH-positive.

### Calcium imaging

Cells were incubated in HEPES-buffered extracellular recording solution (20 mM HEPES, 115 mM NaCl, 5.4 mM KCl, 13.8 mM D-glucose, 1.8 mM CaCl_2_, 0.8 mM MgCl_2_, pH 7.4) containing 1 µM Fura-2 AM (50 µg dissolved in 50 µl dimethylsulfoxide) for 30 min. Cells were then washed and incubated for a further 30 min at room temperature in recording solution before imaging. During recordings, cells were continuously perfused with recording solution (1.8 ml/min) using a peristaltic pump and maintained at 30 °C. All drugs were prepared in recording solution and delivered via this perfusion system.

Prior to recording, cells were illuminated at 568 nm to identify POMC-mCherry-positive cells, and regions of interest (ROIs) were defined, including one background ROI in a cell-free area. During recordings, cells were alternately excited at 340 nm and 380 nm (1 Hz, 50 ms exposure; Xenon arc lamp, Lambda DG-4 Plus), and emitted fluorescence was collected at 510*±*10 nm (CMOS camera, Hamamatsu ORCA-Flash4.0 V3). The fluorescence ratio *r*, used hereafter as the Ca^2+^ signal, was calculated for each ROI as:

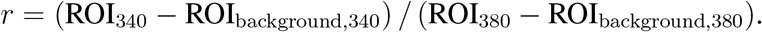

To determine concentration ranges, AVP and CRH were initially tested between 2–10,000 pM. The highest concentrations that did not induce significant desensitisation over three repeated stimulations, using a 20 min inter-stimulation interval, were selected. This yielded final ranges of 2–3000 pM for AVP and 2–2000 pM for CRH.

Calcium imaging experiments comprised a 10 min baseline followed by six 20 min stimulation windows, each with 3 min stimulation and 17 min washout. Cells received five stimulations with AVP alone, CRH alone, or combined CRH+AVP in randomised order to minimise potential systematic order effects, followed by a sixth and final stimulation with 10 mM KCl to assess cell viability.

### Data processing and feature extraction

For each cell, drug-to-cell delivery time (D2C) was determined from the onset of the rapid KCl-evoked Ca^2+^ increase and used as a cell-specific temporal reference. Response time (RT) was determined by first applying a 100 s median filter to the raw trace (`medfilt1`, MAT-LAB; Figure 1A), followed by computation of the approximate derivative (`diff`, MATLAB; Figure 1B). RT was defined as the first time point after D2C at which the derivative exceeded a predefined threshold, calculated as the 25^th^ percentile plus one standard deviation of derivative values between D2C and the peak derivative (Figure 1B). The response window was then defined as the 10 min period following RT (Figure 1C).

**Figure 1:**
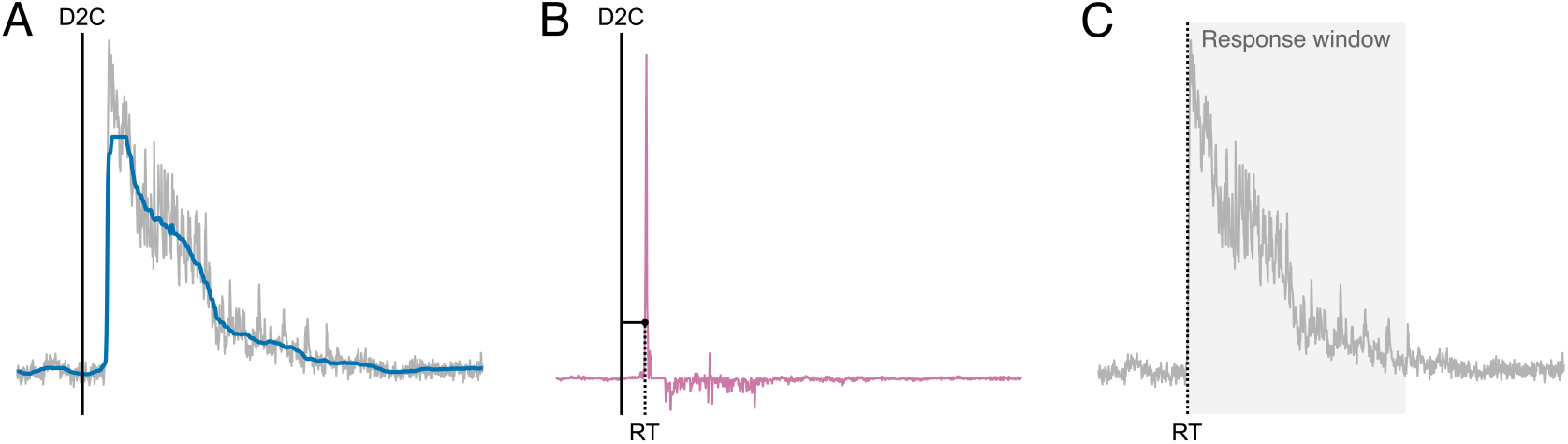
Determination of response time (RT) for individual Ca^2+^ responses. (A) Representative raw (grey) and 100 s median-filtered (blue) Ca^2+^ traces. The black line indicates the drug-to-cell delivery time (D2C). (B) Approximate derivative of the filtered trace (pink), with RT (dashed black line) defined as the first time point after D2C at which the derivative exceeded a predefined threshold (horizontal black line). (C) Raw Ca^2+^ trace with the identified RT and corresponding 10 min response window (grey shaded area).

Because all anterior pituitary cell types were cultured together, and to account for spontaneous Ca^2+^ activity and imperfect viral specificity, we took a conservative approach to cell selection for analysis. Included cells were required to show positive mCherry fluorescence, a robust Ca^2+^ response to 10 mM KCl, and robust Ca^2+^ responses to at least two of the five CRH, AVP, or CRH+AVP stimulations. A robust response was defined as a median Ca^2+^ signal during the first 3 min of the response window that exceeded the baseline median by more than three standard deviations of the baseline signal; baseline was defined as the 3 min pre-RT period.

For each stimulus and each included cell, we computed: (1) the AUC over the response window, calculated from the baseline-adjusted trace, where baseline was defined as the 25^th^ percentile of the signal during the 3 min pre-RT period, with negative AUC values set to zero;(2) the maximum of the Ca^2+^ signal within the response window, measured from the baseline-adjusted trace; and (3) the response delay, defined as the time from D2C to RT.

At the population level, for each stimulation condition and recording, cell recruitment was calculated as the percentage of included cells that showed a robust response.

For each cell and CRH–AVP concentration pair, synergy was quantified by comparing the AUC evoked by co-stimulation with CRH and AVP to the sum of the AUCs evoked by the corresponding individual CRH and AVP stimulations. Synergy was defined as: Synergy = AUC_CRH+AVP_ − (AUC_CRH_ + AUC_AVP_). To account for response heterogeneity, repeated stimulations with 200 pM CRH + 2000 pM AVP were used to estimate a null distribution of absolute synergy values. The 25^th^ percentile of this distribution was 10.87, which was then used to define a null interval of [−10.87, +10.87]. For each CRH–AVP concentration pair, cells with synergy values exceeding +10.87 were classified as *synergistic*, those below −10.87 as *anti-synergistic*, and those within the interval as *non-synergistic*. Cells classified as non-synergistic were excluded from subsequent synergy analyses. For each CRH–AVP concentration pair, the proportions of synergistic and anti-synergistic cells were then calculated across the population.

### Statistical analysis

To evaluate how the features AUC, maximum of the Ca^2+^ signal, and synergy depended on secretagogue concentration, we used linear mixed-effects models. Each feature was modelled as the dependent variable, secretagogue concentration was included as a fixed effect, and random intercepts were included for recording identity nested within cage. Because cells were derived from pooled pituitaries from co-housed animals, cage was used as the biological grouping factor rather than individual animal. Models were implemented in MATLAB using `fitlme` and the formula `feature ∼ secretagogue + (1|cage:rec)`, where `secretagogue` denotes relevant secretagogue concentration(s), and `cage:rec` denotes recording identity nested within cage.

Cell recruitment and the proportions of synergistic and anti-synergistic cells were analysed using generalised linear models with a binomial distribution. The number of recruited, synergistic, or anti-synergistic cells was modelled as the number of successes, with the total number of included cells per recording specified using the `BinomialSize` parameter in MATLAB `fitglm`. Models were specified as `percentage ∼ secretagogue`. Cage-level random effects were evaluated but not retained, as they did not improve model fit relative to the additional model complexity, based on Akaike’s Information Criterion (AIC) and likelihood-ratio tests.

Model slopes, denoted *β*_AVP_ and *β*_CRH_, were used to quantify the overall influence of AVP and CRH concentration, respectively. Secretagogue concentrations were log-transformed before modelling because they spanned several orders of magnitude. AUC, maximum of the Ca^2+^ signal, and synergy values were square-root transformed to better satisfy model assumptions.

Response delay was analysed using a Cox proportional hazards model (`coxph`, R), with time measured from D2C. Robust response onsets were treated as events, whereas cell–stimulus observations without a robust response onset within the 10-min post-D2C detection window were right-censored at 10 min. Secretagogue concentration was included as a fixed effect, and recording identity nested within cage was included as a frailty term, matching the grouping structure used for the linear mixed-effects models. The model was described by the formula `Surv(delay, event) ∼ secretagogue + frailty(cage:rec)`, where `event` is a binary indicator of whether the response to that stimulus was classified as robust. The model coefficients *b*_AVP_ and *b*_CRH_ were exponentiated to obtain hazard ratios, which indicate the relative likelihood of detecting a robust response onset over time.

Non-parametric comparisons between AVP and CRH responses at matched concentrations were performed using Kruskal–Wallis tests.

For all tests, statistical significance was assessed at *p <* 0.05. Analyses were performed in MATLAB R2025b (The MathWorks, Natick, MA, USA) and R version 4.3.1 (R Core Team, 2023) using RStudio 2023.06.1+524.

## Results

### Efficiency and specificity of POMC-mCherry viral targeting

A lentiviral construct expressing mCherry under a POMC promoter selectively targeted corticotrophs in primary pituitary cultures (Figure 2). Among POMC-mCherry-positive cells,82.1 % co-localised with ACTH, whilst 93 % of ACTH-positive cells were mCherry positive, indicating high efficiency and specificity.

**Figure 2:**
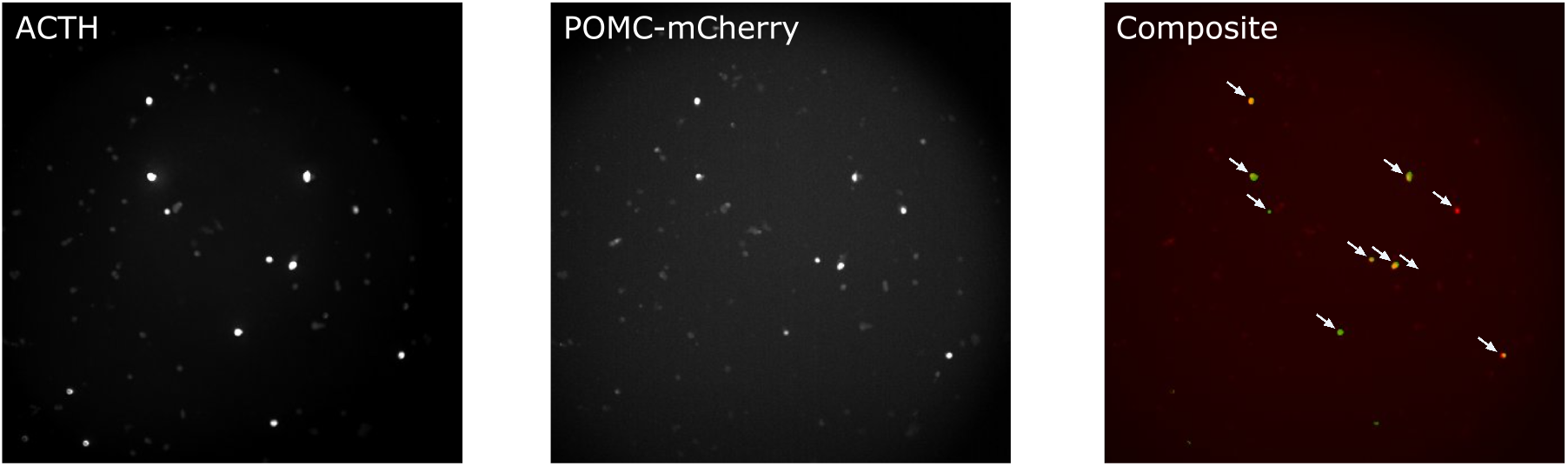
Validation of POMC-mCherry targeting in primary pituitary cells. Cells were transduced with POMC-mCherry lentivirus and fixed 3–4 days later, corresponding to the time of Ca^2+^ recordings, and stained for ACTH. A staining threshold was defined as described in Methods. Regions of co-localisation are indicated by white arrows.

### Concentration-dependent Ca^2+^ responses to AVP and CRH

Stimulation of corticotrophs with AVP or CRH induced rapid increases in the Ca^2+^ signal, although the magnitude, pattern, and duration of responses to a given stimulus were highly heterogeneous across cells (Figure 3). A total of 219 cells met the inclusion criteria for analysis.

**Figure 3:**
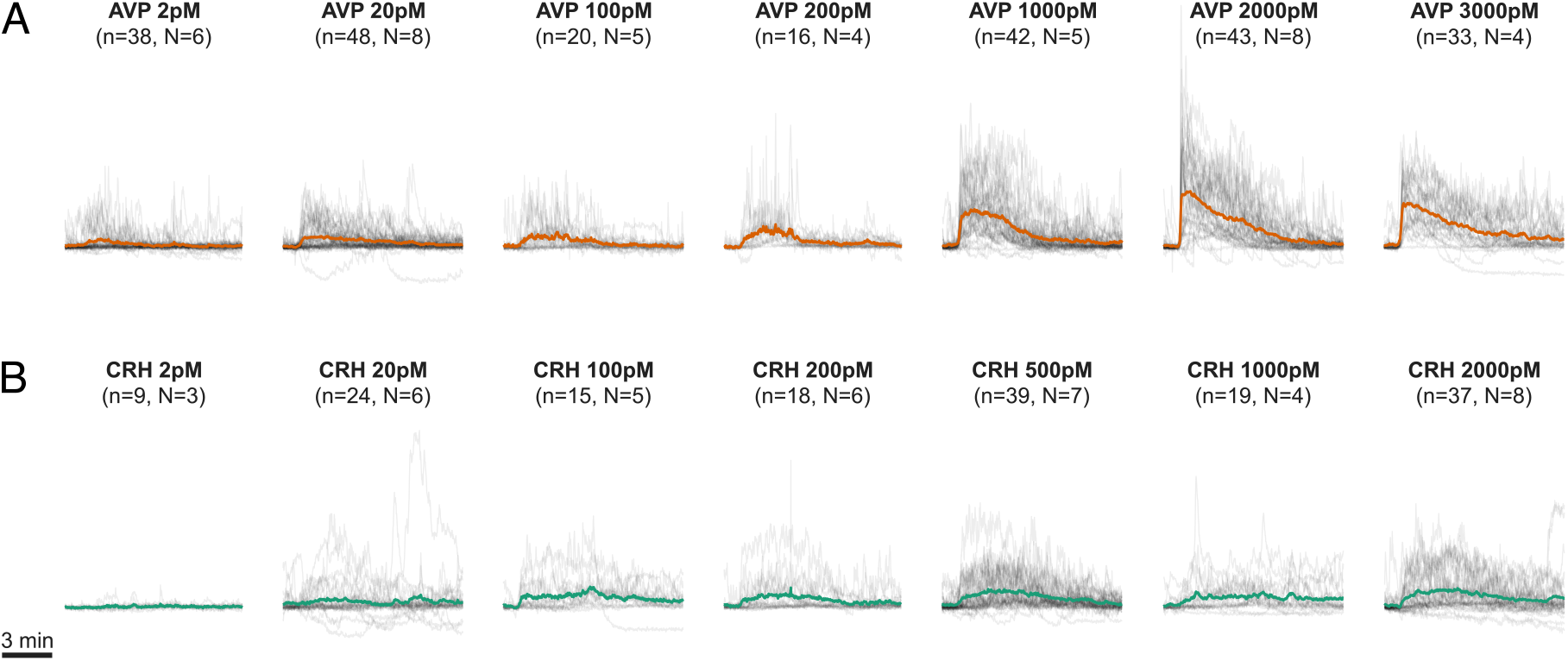
Corticotroph Ca^2+^ responses to 3-min stimulation with AVP (A) or CRH (B). Normalised 340:380 fluorescence ratio traces (*r*) are shown in grey, with mean responses indicated by thick orange (AVP) and green (CRH) lines. The horizontal black bar indicates a 3-min time scale. Traces are aligned to response time (RT). N, number of recordings; n, total number of cells.

AVP and CRH elicited concentration-dependent increases in corticotroph Ca^2+^ responses, reflected in both AUC and the maximum of the Ca^2+^ signal (Figure 4). At matched concentrations, AVP generally evoked larger responses than CRH, with significant differences observed at the concentrations indicated in Figure 4C,F.

**Figure 4:**
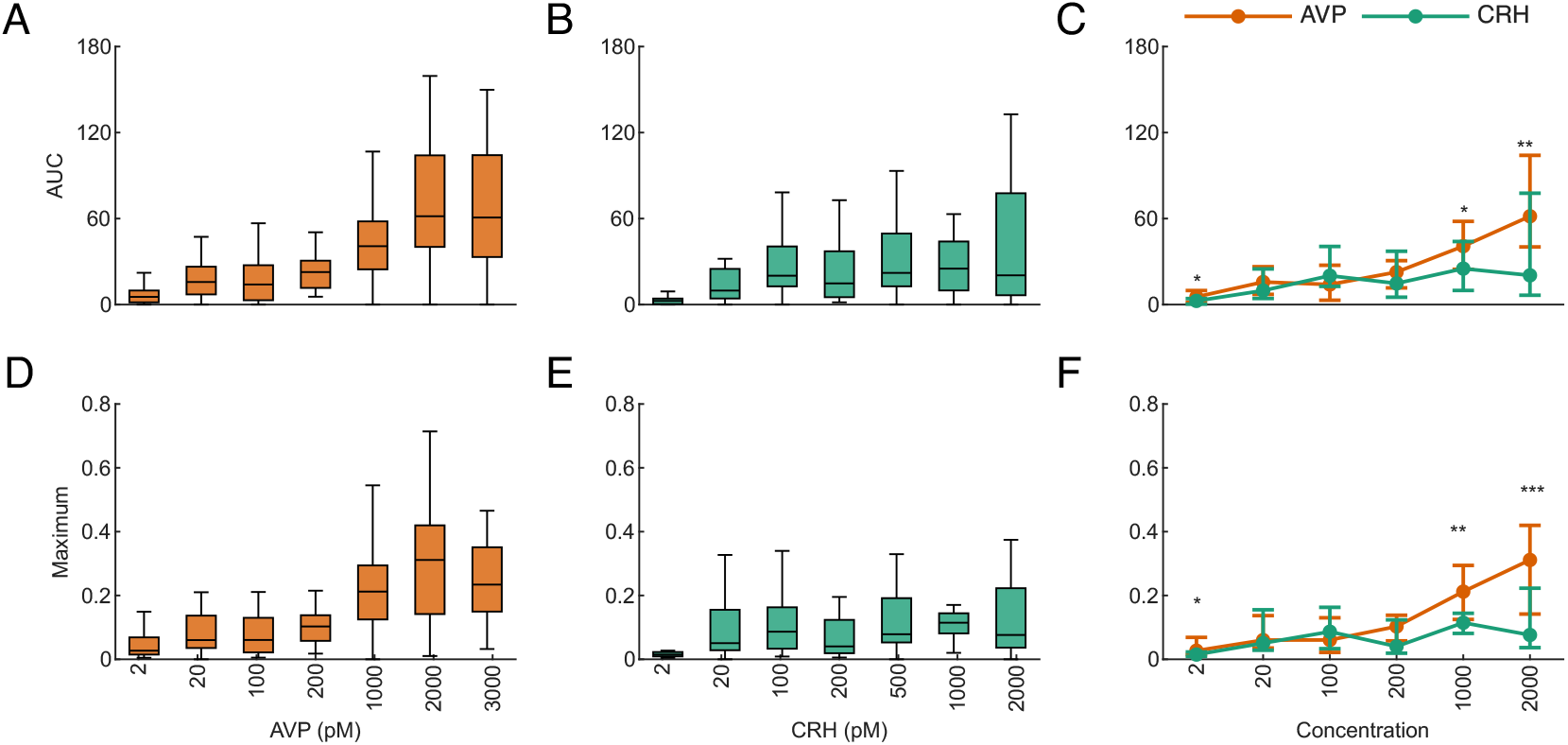
AUC and maximum of the Ca^2+^ signal in corticotroph responses to AVP (orange) and CRH (green). AUC responses to AVP (A) and CRH (B), with comparisons across matched concentrations shown as median (IQR) values (C). Maximum of the Ca^2+^ signal for AVP (D) and CRH (E), with comparisons across matched concentrations shown as median (IQR) values (F). Concentration-dependent effects (A and D; B and E) were assessed using linear mixed-effects modelling. For AVP, both AUC (*β*_AVP_ = 0.728, *p <* 0.001) and maximum of the Ca^2+^ signal (*β*_AVP_ = 0.045, *p <* 0.001) were significantly affected by concentration. Similarly, for CRH, both AUC (*β*_CRH_ = 0.416, *p <* 0.05) and maximum of the Ca^2+^ signal (*β*_CRH_ = 0.020, *p <* 0.01) were significantly affected. Comparisons across matched concentrations (C and F) were performed using the Kruskal–Wallis test. ^*^*p <* 0.05;^**^*p <* 0.01; ^***^*p <* 0.001.

Cell recruitment, calculated as the percentage of included cells showing a robust response, also increased significantly with secretagogue concentration for both AVP and CRH (Figure 5A,B). For CRH, this relationship was sensitive to the inclusion of the 2 pM condition, at which no robust responses were observed: when this concentration was excluded, the effect of CRH no longer showed a significant dependence on concentration (*β*_CRH_ = 0.156, *p >* 0.05), whereas the effect of AVP remained significant (*β*_AVP_ = 0.450, *p <* 0.001). Overall, cell recruitment was higher for AVP than CRH at matched concentrations, reaching statistical significance at the concentrations indicated in Figure 5C.

**Figure 5:**
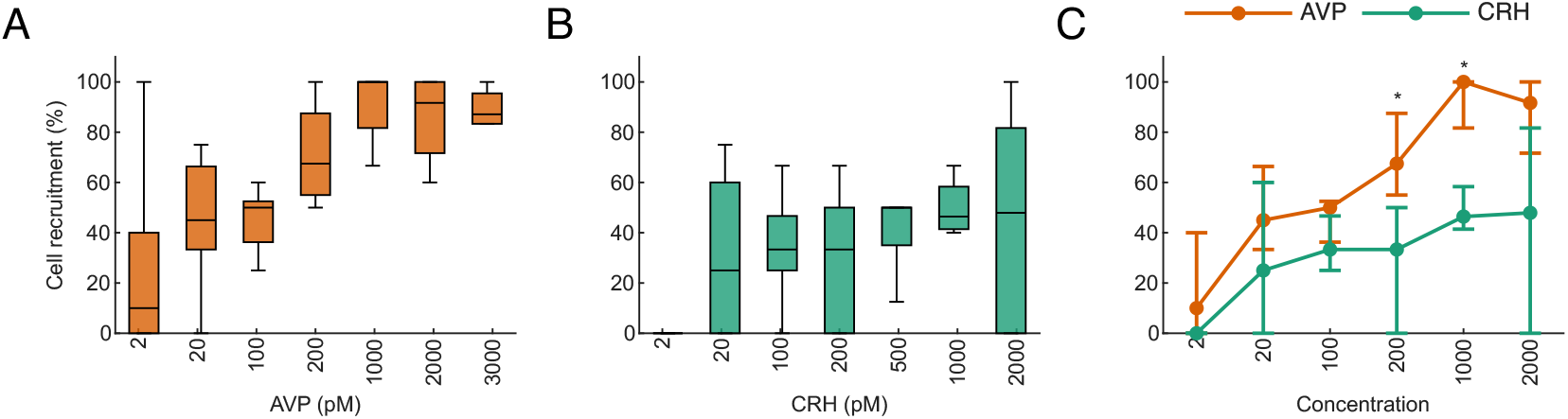
Cell recruitment in response to AVP (A) and CRH (B), with comparisons across matched concentrations shown as median (IQR) values (C). Cell recruitment was calculated for each recording as the percentage of included cells that showed a robust response. No robust responses were detected under stimulation with 2 pM CRH. Concentration-dependent effects were assessed using generalised linear modelling with a binomial distribution. Significant effects were observed for both AVP (*β*_AVP_ = 0.457, *p <* 0.001) and CRH (*β*_CRH_ = 0.251, *p* = 0.008). Comparisons across matched concentrations (C) were performed using the Kruskal–Wallis test. ^*^*p <* 0.05; ^**^*p <* 0.01; ^***^*p <* 0.001.

### Distinct temporal dynamics of corticotroph responses to CRH and AVP

We next assessed response delay, defined as the time from D2C to RT, across AVP and CRH concentrations. Higher concentrations of both AVP and CRH were associated with shorter response delays, consistent with faster activation of Ca^2+^ signalling at higher secretagogue concentrations (Figure 6A,B). At matched concentrations, response delays were generally longer for CRH than AVP at lower concentrations, with significant differences observed at the concentrations indicated in Figure 6C.

**Figure 6:**
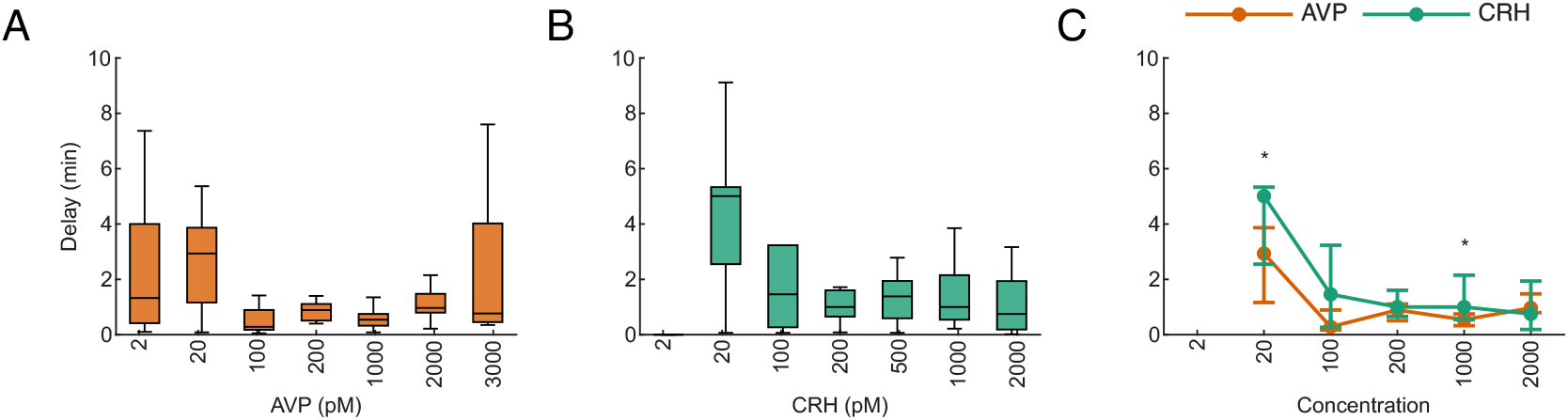
Delay of Ca^2+^ response onsets in corticotrophs to AVP (A) and CRH (B), with comparisons across matched concentrations shown as median (IQR) values (C). Plotted delay values represent observed response onsets; observations without a robust response onset within the detection window were right-censored at 10 min in the Cox model and are not shown. No robust response onsets were detected under stimulation with 2 pM CRH. Concentration-dependent effects on response delay were assessed using Cox proportional hazards regression. Response delay was significantly reduced by increasing AVP (*b*_AVP_ = 0.276, *p <* 0.001) or CRH concentration (*b*_CRH_ = 0.170, *p <* 0.01). Comparisons across matched concentrations (C) were performed using the Kruskal–Wallis test. ^*^*p <* 0.05; ^**^*p <* 0.01;^***^*p <* 0.001.

To further characterise temporal dynamics, we analysed the contribution of each secretagogue to the Ca^2+^ signal over discrete 30 s time windows spanning pre- and post-response periods. Ca^2+^ traces were aligned to RT (black dashed line in Figure 7A,B), and linear mixed-effects modelling was used to estimate time-dependent effects of AVP and CRH. AVP exerted a rapid effect on the Ca^2+^ signal, with *β*_AVP_ increasing sharply around RT and then declining gradually over the post-response period (Figure 7C). Significant effects observed before RT likely reflect subthreshold or gradually developing Ca^2+^ activity that preceded robust response onset. In contrast, the effect of CRH developed more gradually, with *β*_CRH_ increasing after RT, peaking several minutes later, and then declining towards zero (Figure 7D). Together, these analyses indicate that increasing concentrations of both secretagogues accelerate robust response onset, but that AVP and CRH differ in the temporal evolution of the subsequent Ca^2+^ response.

**Figure 7:**
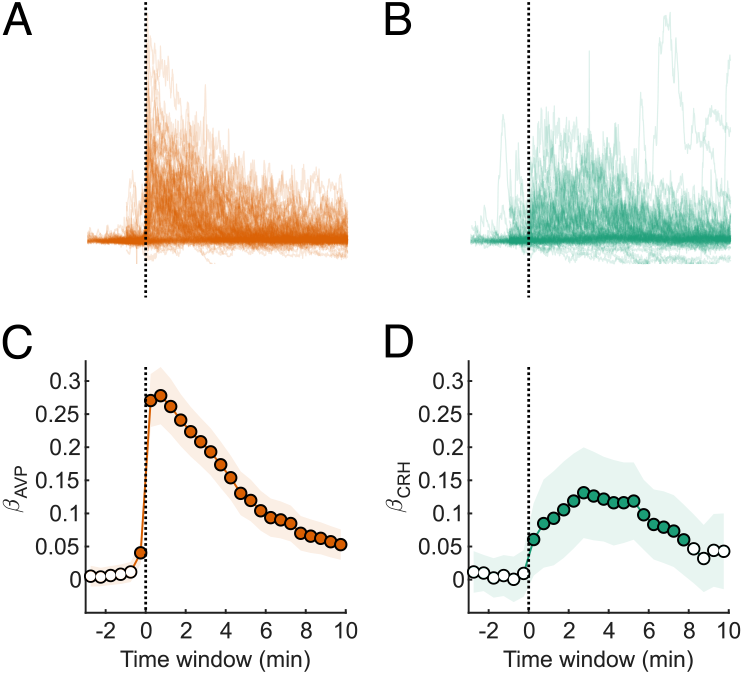
Corticotroph Ca^2+^ responses to 3-minute stimulation with AVP (A) or CRH (B), aligned to response time (black dashed line). Time-dependent effects of AVP (C) and CRH (D) on AUC, estimated across discrete 30-second time windows spanning pre- and post-response periods. Effects (*β*_AVP_ and *β*_CRH_) were estimated using linear mixed-effects modelling. Shaded regions indicate model uncertainty; filled dots denote statistically significant effects; hollow dots indicate non-significance.

### Concentration-dependent interactions between AVP and CRH

AVP and CRH act synergistically to enhance ACTH secretion [24, 25]. To determine whether combined AVP–CRH stimulation alters corticotroph Ca^2+^ signalling, we first assessed responses to co-stimulation by varying the concentration of one secretagogue whilst maintaining the other at a fixed concentration; specifically, AVP was varied in the presence of 200 pM CRH, and CRH was varied in the presence of 2000 pM AVP. Under co-stimulation, increasing AVP concentration in the presence of 200 pM CRH significantly affected AUC, the maximum of the Ca^2+^ signal, response delay, and cell recruitment (Figure 8A,C,E,G). Increasing CRH concentration in the presence of 2000 pM AVP significantly affected AUC and the maximum of the Ca^2+^ signal, but did not significantly affect response delay or cell recruitment (Figure 8B,D,F,H).

**Figure 8:**
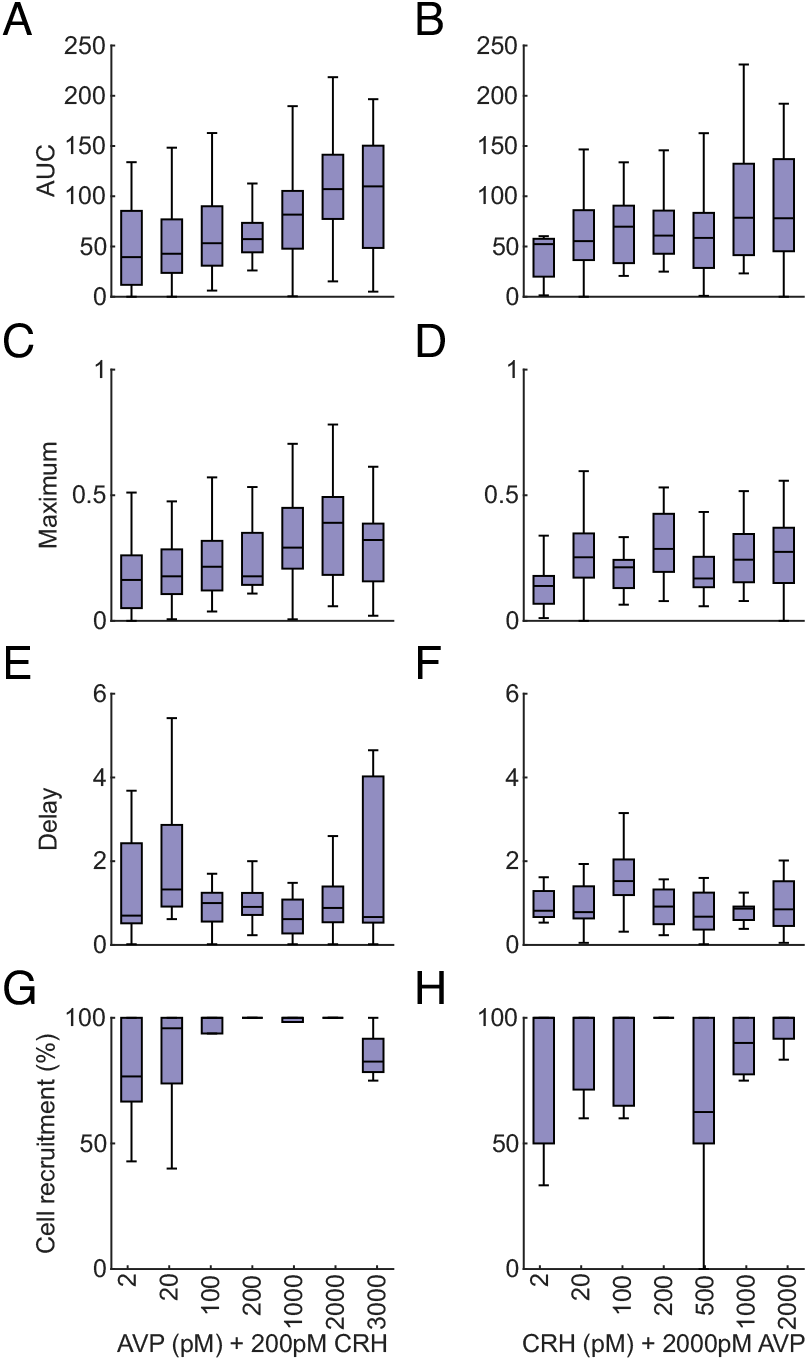
AUC (A and B), maximum of the Ca^2+^ signal (C and D), response delay (E and F), and cell recruitment (G and H) for co-stimulation with AVP and CRH. Experiments in which AVP concentration was varied are shown in the left column, and those in which CRH concentration was varied are shown in the right column. Plotted delay values represent observed response onsets; observations without a robust response onset within the detection window were right-censored at 10 min in the Cox model and are not shown. Concentration-dependent effects were assessed using linear mixed-effects modelling (AUC, maximum of the Ca^2+^ signal), Cox proportional hazards regression (delay), and generalised linear modelling with a binomial distribution (cell recruitment). When AVP concentration was varied in the presence of 200 pM CRH (left column), significant effects were observed for AUC (*β*_AVP_ = 0.538, *p <* 0.001), maximum of the Ca^2+^ signal (*β*_AVP_ = 0.025, *p <* 0.001), response delay (*b*_AVP_ = 0.119, *p <* 0.001), and cell recruitment (*β*_AVP_ = 0.280, *p <* 0.001). When CRH concentration was varied in the presence of 2000 pM AVP (right column), significant effects were observed for AUC (*β*_CRH_ = 0.481, *p <* 0.001) and maximum of the Ca^2+^ signal (*β*_CRH_ = 0.013, *p* = 0.046), but not for response delay (*b*_CRH_ = 0.039, *p >* 0.05) or cell recruitment (*β*_CRH_ = 0.083, *p >* 0.05).

The experimental design included stimulations with AVP alone, CRH alone, and the corresponding AVP+CRH combination, with stimulation order randomised. This allowed synergy to be quantified by comparing the AUC evoked by co-stimulation with the sum of the AUCs evoked by the corresponding individual AVP and CRH stimulations. Representative Ca^2+^ traces from cells stimulated with 2000 pM AVP and 200 pM CRH illustrate the heterogeneous nature of the responses (Figure 9), which were classified as synergistic, non-synergistic, or anti-synergistic according to the criteria described in the Methods.

**Figure 9:**
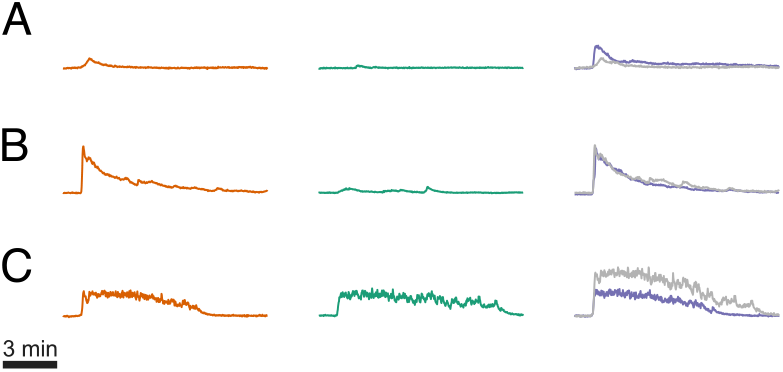
Representative Ca^2+^ traces from three corticotrophs illustrating (A) a synergistic (Synergy = 47.97), (B) a non-synergistic (Synergy =−7.2), and (C) an anti-synergistic (Synergy = −203.73) response to co-stimulation with 2000 pM AVP and 200 pM CRH. Orange, green, and purple traces correspond to responses to 3-minute stimulation with AVP, CRH, and AVP+CRH, respectively. Grey traces represent the sum of responses to AVP and CRH applied in isolation.

For AVP, the proportion of synergistic and anti-synergistic cells did not vary significantly with concentration (Figure 10A,D). In contrast, increasing CRH concentration significantly increased the proportion of synergistic cells without affecting the proportion of anti-synergistic cells (Figure 10B,E). The proportion of synergistic cells differed between AVP and CRH only at the lowest matched concentration (Figure 10C), whilst there were no significant differences between AVP and CRH in the proportion of anti-synergistic cells (Figure 10F).

**Figure 10:**
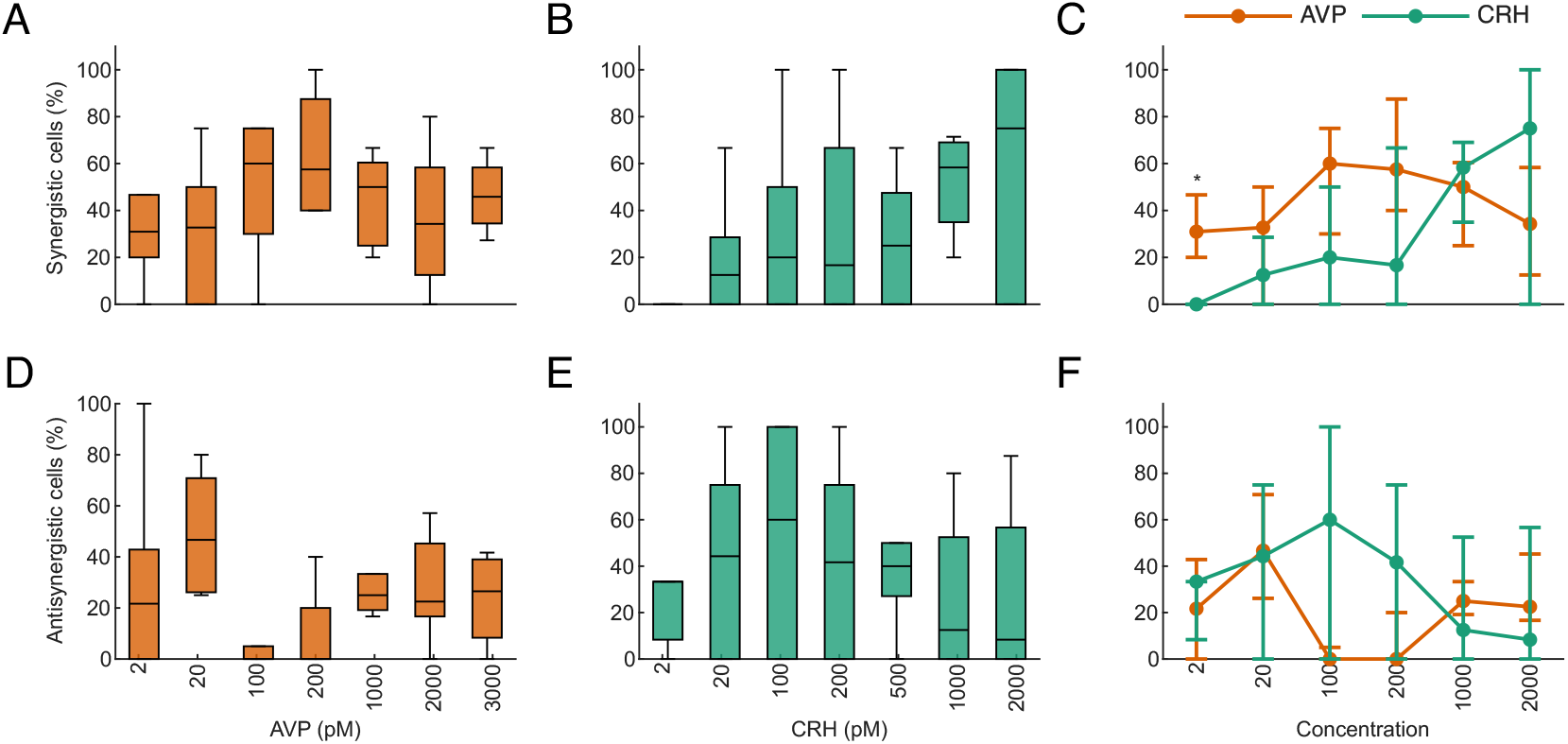
Proportion of synergistic (A–C) and anti-synergistic (D–F) cells across concentrations of AVP (A, D) and CRH (B, E), with comparisons across matched concentrations shown as median (IQR) values (C, F). Experiments in which AVP concentration was varied were performed in the presence of 200 pM CRH, and those in which CRH concentration was varied were performed in the presence of 2000 pM AVP. No synergistic cells were detected under stimulation with 2 pM CRH. Concentration-dependent effects were assessed using generalised linear modelling with a binomial distribution. Changes in AVP concentration did not significantly affect the proportion of synergistic (*β*_AVP_ = 0.022, *p >* 0.05) or anti-synergistic (*β*_AVP_ = −0.021, *p >* 0.05) cells. In contrast, increasing CRH concentration significantly increased the proportion of synergistic cells (*β*_CRH_ = 0.279, *p* = 0.013), but did not significantly affect the proportion of anti-synergistic cells (*β*_CRH_ = −0.019,*p >* 0.05). Comparisons across matched concentrations (C and F) were performed using the Kruskal– Wallis test. ^*^*p <* 0.05; ^**^*p <* 0.01; ^***^*p <* 0.001.

Among cells classified as synergistic, positive synergy values were not significantly affected by either AVP or CRH concentration (Figure 11A,B). Among cells classified as anti-synergistic,negative synergy values were significantly affected by AVP concentration, but not by CRH concentration (Figure 11D,E). No differences in positive or negative synergy values were observed between AVP and CRH at matched concentrations (Figure 11C,F).

**Figure 11:**
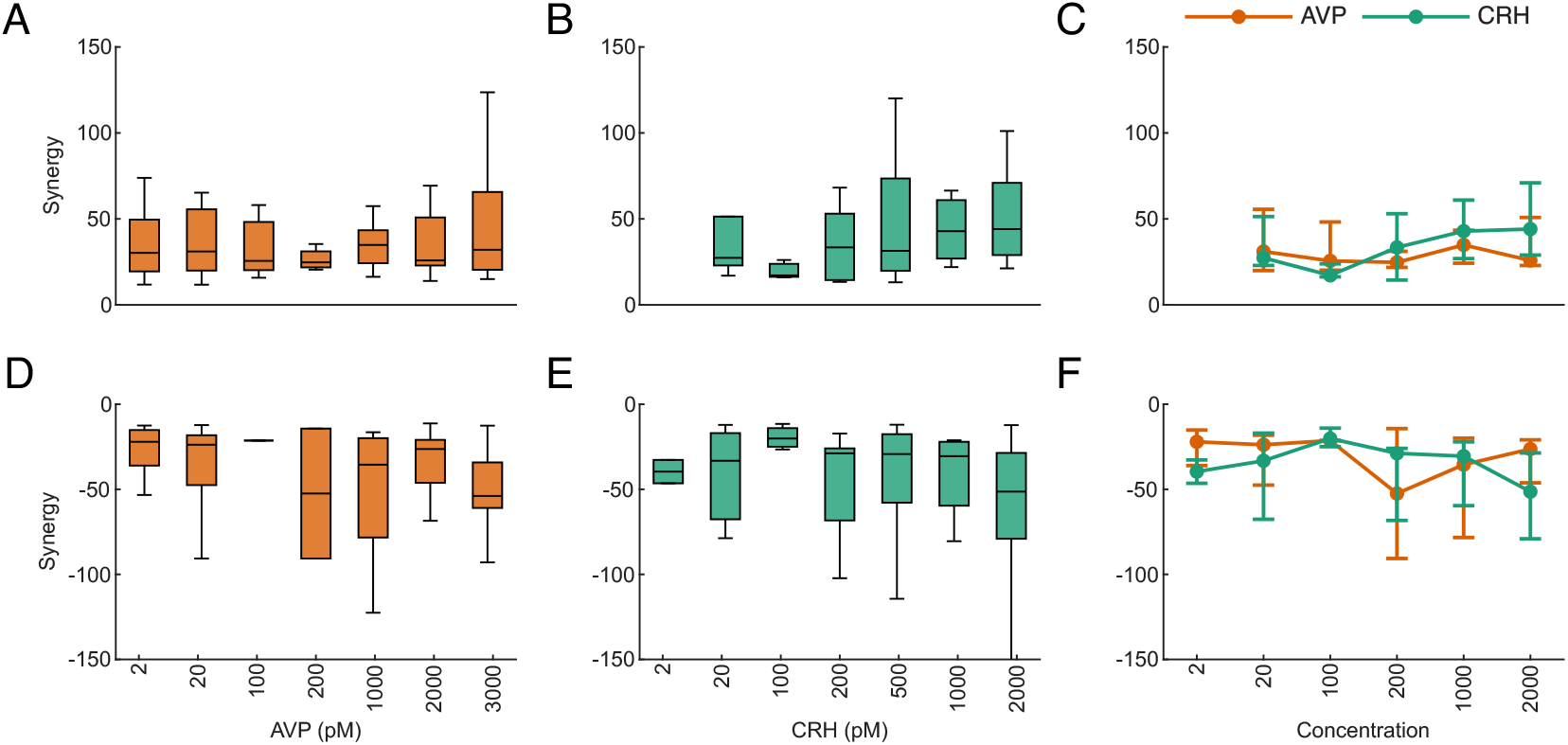
Synergy values across concentrations of AVP (A, D) and CRH (B, E), with comparisons across matched concentrations shown as median (IQR) values (C, F). Panels A–C show positive values classified as synergistic, whereas panels D–F show negative values classified as anti-synergistic. Experiments in which AVP concentration was varied were performed in the presence of 200 pM CRH, and those in which CRH concentration was varied were performed in the presence of 2000 pM AVP. No synergistic cells were detected under stimulation with 2 pM CRH. Concentration-dependent effects were assessed using linear mixed-effects modelling. Changes in AVP concentration did not significantly affect positive synergy values (*β*_AVP_ = 0.050, *p >* 0.05), but did significantly affect negative synergy values (*β*_AVP_ = 0.226, *p* = 0.035). In contrast, increasing CRH concentration did not significantly affect either positive synergy values (*β*_CRH_ = 0.385, *p >* 0.05) or negative synergy values (*β*_CRH_ = 0.165,*p >* 0.05). Comparisons across matched concentrations (C and F) were performed using the Kruskal– Wallis test. ^*^*p <* 0.05; ^**^*p <* 0.01; ^***^*p <* 0.001.

## Discussion

Corticotroph cells of the anterior pituitary are the principal endocrine mediators of hypothalamic stress signals, transducing inputs from parvocellular neurons of the PVN into Ca^2+^-dependent ACTH release. In this study, we investigated how the two primary stress-related neurohormones, CRH and AVP, shape intracellular Ca^2+^signalling in isolated corticotroph cells when applied individually and in combination across physiologically relevant concentration ranges. By examining response magnitude, temporal dynamics, and cell recruitment, this study provides insight into how corticotroph activity is regulated across different modes of HPA axis activation.

In vivo studies have implicated CRH as a key regulator of basal ultradian corticosterone rhythms [12], with AVP appearing to contribute more prominently during chronic or repeated stress conditions [26, 27, 28]. This context-dependent shift is supported by increased AVP gene and protein expression in PVN neurons under sustained stress conditions [29, 30], and by evidence that AVP helps maintain HPA axis drive when CRH signalling is downregulated [31, 28, 27]. We found that AVP evoked larger Ca^2+^ responses than CRH at several matched concentrations, spanning the lower and upper ends of the tested range. These findings suggest that AVP’s functional relevance extends beyond chronic stress states, where AVP levels may be elevated, and that AVP can act as a potent regulator of corticotroph Ca^2+^ signalling across a broader range of secretagogue concentrations.

Analysis of cell recruitment supported a greater recruitment capacity for AVP than CRH. AVP drove robust cell recruitment across its concentration range, with the highest concentrations recruiting most of the analysed corticotroph population. Although CRH also showed an apparent concentration-dependent increase in cell recruitment, this relationship was strongly influenced by the absence of robust responses at 2 pM CRH and was no longer significant when this concentration was excluded. Moreover, even at the highest concentration tested, CRH recruited a smaller fraction of cells than AVP. Together, these findings suggest that AVP is capable of recruiting a larger fraction of the corticotroph population than CRH across the tested concentration range.

One possible explanation for the greater corticotroph recruitment by AVP, particularly at the highest tested concentrations, is that CRH and AVP engage partly distinct Ca^2+^ signalling mechanisms. CRH-evoked responses may rely more strongly on membrane excitability and calcium influx through voltage-dependent channels, whereas AVP can also mobilise calcium from intracellular stores through phospholipase C-linked signalling. Cells that are less electrically excitable may therefore be less likely to generate a robust CRH response, whilst remaining capable of responding to AVP. In addition, although CRHR1 expression has been demonstrated in corticotrophs [32, 33], CRHR1 expression may not be uniform across the corticotroph population, and receptor expression alone may not be sufficient to confer robust Ca^2+^ responsivity in all cells. Intercellular variability in receptor abundance, downstream signalling capacity, or technical limitations in the detection of low-amplitude Ca^2+^ responses may therefore contribute to the observed variability in cell recruitment.

Marked heterogeneity in corticotroph Ca^2+^ signalling was observed across AUC, the maximum of the Ca^2+^ signal, response delay, and cell recruitment. Whilst heterogeneous corticotroph Ca^2+^ signalling has been reported previously [21], this study is, to our knowledge, the first to examine how such variability changes with AVP and CRH concentration. Functional heterogeneity can expand response dynamic range, enhance sensitivity to graded inputs, and support stimulus encoding, and has been observed in other pituitary cell types, including gonadotrophs [34], lactotrophs [35], and somatotrophs [36]. In this context, heterogeneity across secretagogue concentrations may help corticotroph populations encode graded hypothalamic inputs whilst maintaining responsiveness across different levels of HPA axis activation.

The graded nature of cell recruitment across secretagogue concentrations further suggests corticotroph subpopulations with distinct activation thresholds. Previous functional, morphological, and transcriptomic studies support the existence of corticotroph heterogeneity within the anterior pituitary [37, 38, 39, 40]. Although our experiments were not designed to define morphological or transcriptional populations, the observation that some corticotrophs were recruited only at higher secretagogue concentrations, whereas others remained unresponsive, is consistent with heterogeneous activation thresholds within the corticotroph population.

Temporal analysis revealed distinct dynamics of AVP- and CRH-evoked Ca^2+^ responses. Increasing concentrations of both secretagogues reduced response delay, indicating more rapid activation of Ca^2+^ signalling at higher concentrations, although CRH responses were generally slower than AVP responses at the lower end of the tested range. Time-resolved modelling showed that the AVP effect emerged rapidly around RT and then declined gradually across the post-response period. In contrast, the CRH effect developed more slowly, peaked several minutes after RT, and then declined. These temporal profiles may reflect distinct downstream signalling mechanisms, with CRH acting primarily through CRHR1-mediated regulation of membrane excitability and voltage-gated calcium influx [41], and AVP acting through V1b receptors to mobilise calcium from intracellular stores [42]. The persistence of the AVP-associated effect may also reflect secondary activation of voltage-gated calcium channels [43]. Future studies using voltage-gated calcium channel inhibition or intracellular-store depletion could help define the relative contribution of these pathways to AVP- and CRH-evoked Ca^2+^ responses.

Synergistic interactions between CRH and AVP in driving ACTH secretion have been well documented in vivo, with early studies demonstrating that AVP markedly potentiates the corticotrophic actions of CRH [25, 44]. Subsets of parvocellular PVN neurons also package CRH and AVP within the same median eminence neurosecretory vesicles [45], making simultaneous release likely under physiological conditions. To examine how such convergent inputs are integrated at the level of intracellular Ca^2+^ signalling, we analysed responses to combined CRH–AVP stimulation. Under co-stimulation, AUC and the maximum of the Ca^2+^ signal remained concentration-dependent for both secretagogues. However, response delay and cell recruitment were concentration-dependent only when AVP was varied in the presence of CRH, and not when CRH was varied in the presence of AVP. This suggests that, under co-stimulation, AVP remains a stronger determinant of response timing and recruitment, whereas both secretagogues continue to shape response magnitude.

Increasing CRH concentration significantly increased the proportion of corticotrophs classified as synergistic, without affecting the proportion classified as anti-synergistic. In contrast, varying AVP concentration did not alter the proportion of synergistic or anti-synergistic cells. Positive synergy values were not significantly affected by either secretagogue, suggesting that CRH concentration influenced the likelihood that a cell exhibited synergy, rather than the magnitude of positive synergy once present. Negative synergy values were affected by AVP concentration, indicating that anti-synergistic interactions may be regulated differently from synergistic interactions. These analyses are confined to intracellular Ca^2+^ responses, and synergistic effects may be further amplified at the level of ACTH secretion, which was not assessed here.

Although CRH alone evoked smaller Ca^2+^ responses and recruited fewer corticotrophs than AVP, its ability to increase the proportion of synergistic cells under co-stimulation suggests an important role in signal integration. Thus, CRH may exert substantial influence on pituitary output not by recruiting the largest population of corticotrophs when acting alone, but by increasing the likelihood that cells respond synergistically when AVP is also present. Future studies using single-cell secretion assays will be required to determine how calcium-level synergy maps onto ACTH release and to resolve the functional consequences of CRH–AVP co-stimulation.

Ultimately, this study demonstrates that hypothalamic secretagogue concentration shapes corticotroph Ca^2+^ signalling across multiple response features. AVP generally evoked larger responses, stronger cell recruitment, and faster response onset than CRH, whereas CRH selectively increased the proportion of cells classified as synergistic under co-stimulation. Together, these findings highlight functional heterogeneity as a core feature of corticotroph Ca^2+^ signalling and suggest that AVP and CRH make distinct, concentration-dependent contributions to corticotroph activation.

This work used a dissociated single-cell model, enabling resolution of individual cell behaviour but lacking the spatial organisation, paracrine interactions, and physiological feedback present in vivo. Future studies using more physiologically representative preparations, such as pituitary slice models, will help define how these mechanisms operate within the intact system. Despite these limitations, our findings provide insight into how hypothalamic inputs of differing concentration and combination may interact to fine-tune corticotroph Ca^2+^ signalling.

## Acknowledgements

We thank Heather McClafferty and Mike Shipston (University of Edinburgh, United Kingdom) for providing the original POMC reporter construct used to generate the lentiviral construct in this study.

